# Physical and functional interactome atlas of human receptor tyrosine kinases

**DOI:** 10.1101/2021.09.17.460748

**Authors:** Kari Salokas, Tiina Öhman, Xiaonan Liu, Iftekhar Chowdhury, Lisa Gawriyski, Salla Keskitalo, Markku Varjosalo

## Abstract

Much cell-to-cell communication is facilitated by cell surface receptor tyrosine kinases (RTKs). These proteins phosphorylate their downstream cytoplasmic substrates in response to stimuli such as growth factors. Despite their central roles, the functions of many RTKs are still poorly understood. To resolve the lack of systematic knowledge, we used three complementary methods to map the molecular context and substrate profiles of RTKs. We used affinity purification coupled to mass spectrometry (AP-MS) to characterize stable binding partners and RTK-protein complexes, proximity-dependent biotin identification (BioID) to identify transient and proximal interactions, and an *in vitro* kinase assay to identify RTK substrates. To identify how kinase interactions depend on kinase activity, we also used kinase-deficient mutants. Our data represent a comprehensive, systemic mapping of RTK interactions and substrates. This resource adds information regarding well-studied RTKs, offers insights into the functions of less well-studied RTKs, and highlights RTK-RTK interactions and shared signaling pathways.

## Introduction

Protein phosphorylation reversibly controls the activity or localization of many proteins and is dynamically regulated by protein kinases and protein phosphatases, which phosphorylate and dephosphorylate proteins, respectively. Protein kinases catalyze the transfer of a phosphate group from ATP to threonine, serine, and tyrosine amino acids of specific target proteins. Currently, 571 human protein kinases have been identified. Of these, 137 are tyrosine kinases. Receptor tyrosine kinases (RTKs) are a subclass of tyrosine kinases that act as initiators, amplifiers, and central nodes in a plethora of complex biological functions and are mainly associated with intercellular communication. RTKs regulate key properties of their substrate proteins, which is essential for the coordinated actions of biological pathways and processes. Similar to other kinases, RTKs are strongly associated with a multitude of human diseases, such as cancer and a variety of multifactorial diseases and developmental disorders (McDonell et al., 2015).

In the human genome, 58 RTKs have been identified (Robinson et al., 2000). These RTKs are classified into 20 different subfamilies containing between 1 and 14 members. The Ephrin receptor subfamily is the largest, with 14 members (Liang et al., 2019; Pasquale, 2005), followed by the PDGF subfamily, which includes 5 RTKs (Demoulin and Essaghir, 2014; Kazlauskas, 2017), and the ErbB (Hynes and MacDonald, 2009; Warren and Landgraf, 2006) and FGF groups (Goetz and Mohammadi, 2013; Turner and Grose, 2010), each with four members. The other subfamilies have three or fewer members. While some RTKs, such as EGFR or ERBB2 (also known as HER2), have been extensively studied, most RTKs have been less well studied and have few known interactors; consequently, our understanding of their substrates of protein-protein interaction partners is quite limited.

RTKs are thought to exist on the cell membrane as monomers, dimers, and oligomers. While dimerization or oligomerization is required for activation (Lemmon and Schlessinger, 2010), not all dimers or oligomers actively signal (Clayton et al., 2005; Gadella and Jovin, 1995; Ward et al., 2007). Once oligomerization has occurred, the intracellular domains can transphosphorylate one or more tyrosine in neighboring RTKs. In addition to canonical cell surface signaling, nuclear signaling activity has also been identified for multiple RTKs (Song et al., 2013). The phosphorylated receptor serves as a platform for the assembly and activation of intracellular signaling intermediaries. An inactive kinase is in an autoinhibitory conformation, and this conformation is released by the phosphorylation of an activation loop, after which signaling can proceed. Protein kinases are kept inactive by phosphatases. Protein tyrosine phosphatases (PTPs), in addition to deactivating RTKs when appropriate, also function to maintain RTKs in an inactive state. Indeed, inducing the activation of RTKs is possible in one of two ways: ligand binding or inhibition by cellular phosphatases (Ostman and Böhmer, 2001; Reynolds et al., 2003; Tonks, 2006). PTPs, in turn, can be inhibited *in vitro* with vanadate or pervanadate, leading to tyrosine kinase activation (Boersema et al., 2010; Huyer et al., 1997; Zhao et al., 1996).

RTKs exert changes via interactions with other proteins and by phosphorylating their substrate proteins. The interactions can be stable, as in the case of stable protein complexes, or they can be short-lived transient associations. Therefore, to understand the role of RTKs in cellular signaling networks, it is vital to map their protein-protein interaction (PPI) networks. This goal, however, is hindered because a large number of RTKs have few known interactors. Two well-established and reliable methods for mapping PPIs by mass spectrometry are affinity purification coupled to mass spectrometry (AP-MS) and proximity-dependent biotin identification (BioID). AP-MS captures stable interactions and can quantitatively capture other complex components in addition to direct interactors. BioID, in contrast, does not require a stable interaction but can also capture transient interactions within an ∼10 nm radius. Multiple proteins may be identified with multiple baits, which suggests that these proteins participate in the same process or protein complex (Drew et al., 2017; Knight et al., 2017; Youn et al., 2018).

In this study, we performed systematic AP-MS and BioID analyses of ∼90% of human RTKs in their activated state. This set of 52 RTKs included 7 RTKs with fewer than 20 previously identified interactors. The generated interactome network included > 6000 unique high-confidence RTK-protein interactions. Furthermore, to detect interactions that depended on the corresponding kinase activity, we used kinase activity-deficient (KD) mutants for 11 RTKs. Additionally, we used a phosphoproteomic approach to identify substrates for 45 RTKs. The results represent a comprehensive RTK interaction network and reveal central pathways through which RTKs may exert their effects, as well as networks of probable associations between interactor proteins and RTK-specific functional enrichment.

## Results

### Defining the RTK interaction landscape

To comprehensively identify RTK-interacting proteins, we used two complementary methods, AP-MS and BioID MS. First, 52 human RTKs were cloned into the MAC-tagged expression vector (Liu et al., 2018) and inducibly expressed in 52 stable cell lines. In all cases, the C-terminal tag was used. Each of these cell lines had the corresponding MAC-tagged RTK incorporated in a single genomic locus, and expression could be induced. AP-MS allows the capture of stable interactions and the derivation of complex stoichiometry, while BioID can also detect proximal and transient interactions (Liu et al., 2018) (Fig. S1A). To capture the interactions of active RTKs, cellular PTPs were inhibited with pervanadate prior to sample collection. Pervanadate irreversibly inhibits PTPs by modifying the catalytic cysteine (Huyer et al., 1997).

The 52 RTKs (>90% of all human RTKs) studied here include all RTK subfamilies (Fig. 1A) (Lemmon and Schlessinger, 2010). After stringent statistical filtering, we identified 6050 unique high-confidence interactors (HCIs) (Table S1A). A total of 1145 interactions were identified with AP-MS, 4497 with BioID, and 408 with both methods. The interactors consisted of 1521 unique proteins. The number of identified interactors varied significantly between individual kinases, but many RTK subfamilies showed similar numbers of interactors. The number of known interactions identified was significantly higher than what would be expected from random interaction networks with the same topology as the RTK network (Fig. 1A, inset). The information gathered in this study therefore supplements the scarce interaction data available for many less well-studied RTKs.

**Figure 1.**
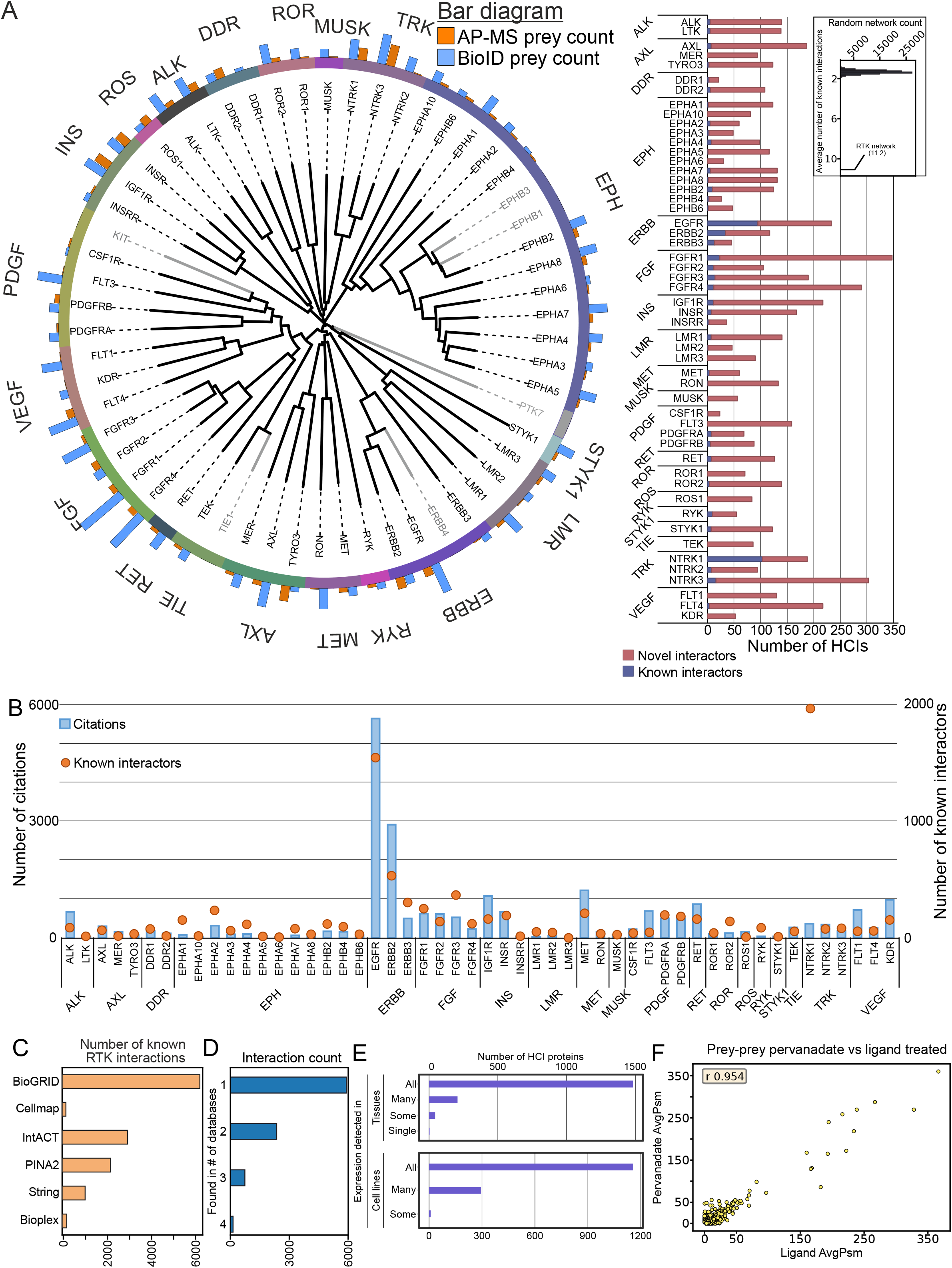
General assessment of study scope and interaction data landscape. A. Left: Sequence alignment tree of the receptor tyrosine kinase (RTK) family. Members of the 20 different receptor tyrosine kinase subfamilies are grouped according to their sequence (kinase domain) homology to their respective subfamilies, indicated by the unique colors. Gray color indicates RTKs not included in this study. Number of high-confidence interactor (HCI) proteins identified in AP-MS (orange) and BioID (blue) experiments are indicated above the circle. Right: Comparison of the detected interactions to existing knowledge. The number of HCIs detected in this study are divided to previously reported interactions (blue) and novel interactions (red). The number of known interactors was highly enriched compared to random networks of the same topology (inset). B. Number of citations and known interactors per RTK, which are grouped into their respective subfamilies. Citations are shown in blue bars and plotted against the Y-axis on the left, while known interactors are shown with orange bubbles, and the right axis. C. Number of known RTK interactors from each of the databases used for the known set. D. Number of known RTK interactors seen in one or more of the used databases. E. Expression of identified HCI proteins in tissues (top) and cell lines (bottom) from human protein atlas (Uhlén et al., 2015). Detected in all: expression detected in all available tissues or cell lines; detected in many: detected in at least a third of the tissues/cell lines; detected in some: detected in more than one, but fewer than a third of the tissues/cell lines. F. HCI comparison between ligand- and pervanadate-treated samples for 8 RTKs.

15 RTKs had more than 150 identified interactors, and the remaining 37 had fewer interactors (Fig. 1A). While some RTKs have been well-studied with many known interactions, most have only a few reported interactions (Fig. 1B), highlighting the need for a systematic study. The number of known interactors for RTKs generally follows the number of citations for each RTK (Fig. 1B), and indeed 19 RTKs had fewer than 100 publications associated with them in the NCBI publication database. For known interactions, we utilized a database combining six databases of interactions (Fig. 1C).

We next decided to characterize the distribution of the known interactors across the six (6) databases from which they were taken. BioGRID, IntACT, and PINA2 contributed the highest number, followed by String, and finally bioplex and human cell map. To characterize how commonly seen the known interactors were, we next analyzed how many databases each interaction was featured in (Fig. 1D). While most interactions were only seen in one database, roughly a third of the interactions were shared between two or more. The largest proportion were seen at least two databases as expected, considering the complimentary nature of BioGRID, IntAct, and PINA2.

Many RTKs share interactions with members of their own subfamily (Fig. S1B). While most subfamilies have a high degree of interconnected interactors, each RTK in this study has identified HCIs, which were not shown to interact with other members of their respective subfamilies. For example, the Eph subfamily has many shared interactions, while the ERBB, INS, and LMR subfamilies have fewer shared interactions, which may indicate similar functions within the Eph family. A second source of variability is the interaction types themselves. BioID interactions represent a higher proportion of all interactions in all subfamilies, except for ROS. However, in different subfamilies, the proportion of BioID interactions varied from 87% with VEGF to 40% with ROS. Shared interactors were often identified with both methods (e.g., the shared cluster in the ERBB subfamily): 27% of the interactions shared between receptors in the same subfamily were detected with both methods, whereas 15% of interactions overall were detected with both methods. The higher percentage may suggest the presence of proteins that are instrumental to the overlapping functions of the receptors in the subfamily. Interactors were widely shared across subfamily boundaries as well. We detected 675 interactors shared within subfamilies and 728 shared with receptors in another subfamily (Fig. S1C, Table S1A). Common HCIs may suggest potential RTK functional overlap and crosstalk, while unique HCIs may indicate receptor-specific functions and RTK-specific variations in possible shared pathways.

To determine whether we could identify indications of the active state of the bait RTKs, we analyzed the AP-MS data for known autophosphorylation site(s) for each RTK. For the majority of RTKs, we identified known tyrosine autophosphorylation site(s) as phosphorylated site(s) (Table S2). In order to further validate the phosphorylation status of the bait RTKs, we performed an anti-phosphotyrosine western blot (WB) analysis of a subset of the RTKs (Fig. S1D), and detected phosphorylation in all of the 8 RTKs analyzed. To ensure that MAC-tagged RTKs localize to plasma membrane, we carried out immunofluorescence confocal microscopy imaging for all of the baits included in the study (Fig. S2).

Given the varied expression of RTKs across tissues and cell types, we also decided to analyze, whether the interactions detected could be cell-line specific, or proteins that are expressed in a variety of tissues. For this purpose, we mapped expression level data from the human protein atlas (Uhlén et al., 2015) project (Fig. 1E). We next divided the identified interactors based on annotations of the database into proteins that were detected in all, many (>=33 %), some (>1) or one cell line or tissue type. The majority of our unique interactors were seen across all tissues and cell lines included in the atlas, while fewer than 300 were seen in many, and fewer than 100 in some or only one.

We next utilized a subset of RTKs to investigate the effect of pervanadate treatment in comparison to ligand-induced activation. We performed side by side AP-MS and BioID experiments with pervanadate-treated, and ligand-treated cell lines of 8 RTKs (EGFR, FGFR1, FGFR4, IGF1R, INSR, INSRR, PDGFRB, and RET). For these RTKs the main ligand was known, and they were available as recombinant protein with validated activity. From these experiments, we identified in total 1132 high-confidence interactions, consisting of 595 unique proteins. Of these, ∼ 80% (872) of the HCIs were seen in both pervanadate and ligand-treated cells. The majority of the prey proteins were seen with similar spectral count values in both experiments (correlation value 0.954, Fig. 1F). Of the interactions seen only in ligand-treated samples, 83 were detected with an average spectral count of over 5. Of these, 61 were seen only in AP-MS experiments, 18 in BioID, and four in both. Likewise, 25 HCIs were seen only in pervanadate-treated samples (14 AP-MS only, 10 BioID, and one in both). On the functional level, however, the proteins which were seen only in either pervanadate- or ligand-treated experiments fell into the same functional groups with proteins that were identified in both experiments (Table S1C).

### Kinase-kinase interactions between RTKs

To investigate whether RTK heterodimers or -oligomers contributed to the number of identified shared HCIs, we next investigated the presence of RTK-RTK interactions in detail (Fig. 2A). In total, we identified 77 RTK-RTK interactions, of which 33 were between receptors in the same subfamily. The majority of these subfamily interactions (27) were detected either with AP-MS or both AP-MS and BioID. In contrast, 27 of the 44 interactions between receptors in different subfamilies were detected via BioID only. The identifications derived from BioID alone could more specifically indicate membrane areas and structures commonly shared between the RTKs than identifications derived by other methods. However, the 16 RTK-RTK interactions that were detected by both methods and 28 detected via AP-MS alone suggest the formation of a wide variety of stable RTK-RTK heterodimers. While heterodimerization is a well-documented phenomenon in RTKs, many of the specific interactions here have not been documented previously. Eighteen of the 77 (23%) were previously known, leaving 59 (77%) novel interactions. To validate the RTK-RTK interactions, we performed co-IP analysis of 27 RTK-RTK interactions that were seen in AP-MS data, and detected the interactions with all but 4 of them (Fig. S1E), possibly indicating that these four interactions are not direct but mediated by another protein.

**Figure 2.**
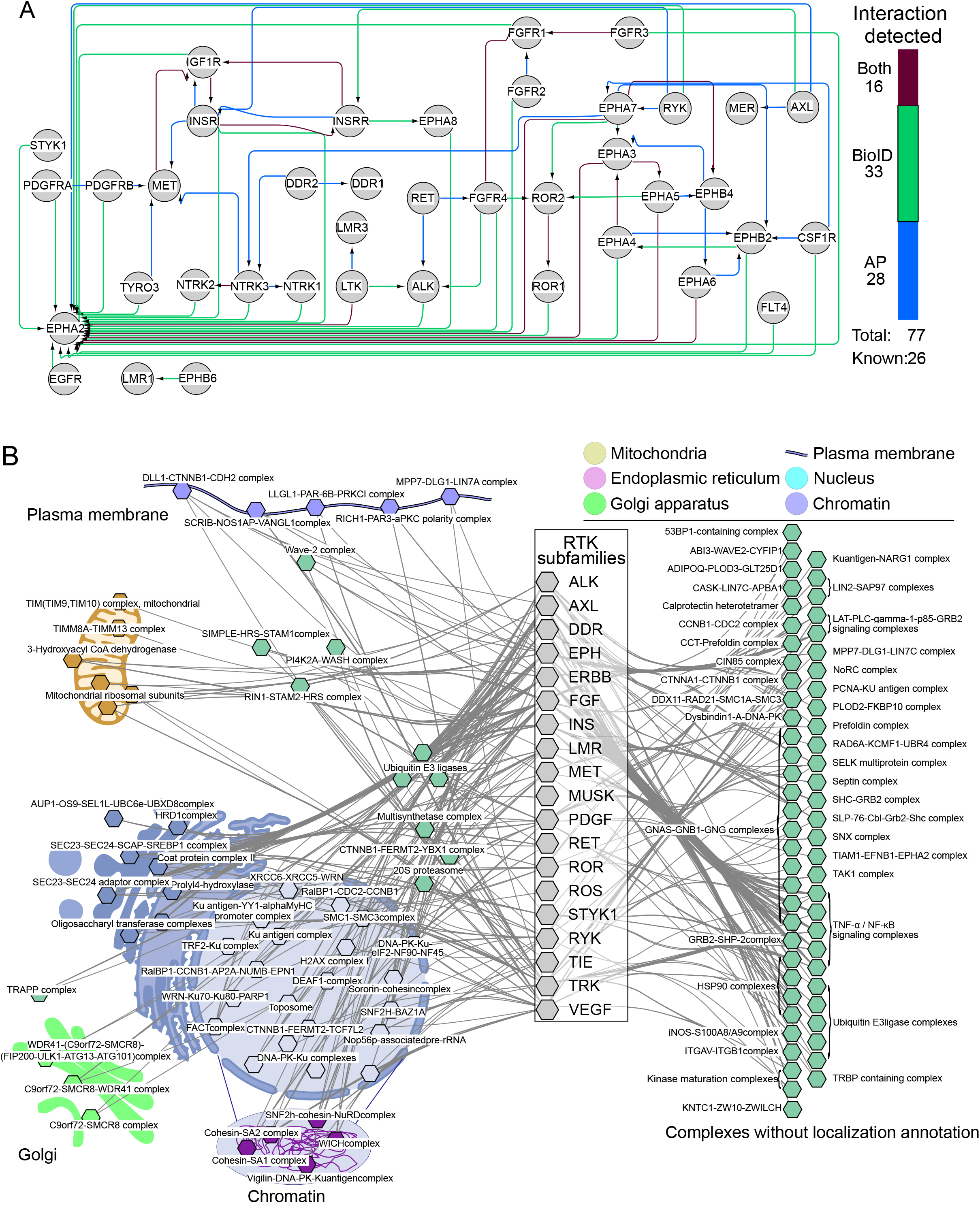
Bait-bait interactions and enriched complexes in the RTK data. A. High-confidence bait-bait interactions detected between the RTKs. Connections are colored based on whether they were detected in AP-MS (blue), BioID (green) or both (burgundy). B. Significantly (q<0.05) enriched CORUM complexes in the interactomes of the RTK subfamilies. The cellular localization was assigned to each complex with available GO cellular component in CORUM. Connections from subfamilies to complexes denote significant enrichment of the complex with one or more members of the subfamily. On the right side, complexes without localization information are grouped based on their protein composition.

Interestingly, EphA2 was seen with a majority of RTKs (30 in total), although it was previously known to form complexes only with EGFR, ErbB2, EphA7, DDR1, and NTRK3 (Brantley-Sieders et al., 2008; De Robertis et al., 2017; Huttlin et al., 2017; Larsen et al., 2007; Lemeer et al., 2012; Oricchio et al., 2011; Zhuang et al., 2010). Through AP-MS only or both methods, we detected six interactions between EphA2 and another RTK (AXL, EphA3, EphA5, EphA6, EphA7, and LTK). Of these, only EphA7 is a previously known interactor. To our knowledge, EphA2 is not highly expressed in HEK-293 cells (Table S1B); hence, its wide identification is unlikely to be due to expression levels. It may therefore be possible that the identified AP-MS interactions of RTKs with EphA2 represent heterocomplexes, while proximal or transient interactions may be due to localization with similar membrane and internalization compartments.

### RTK interactors participate in complexes in a wide variety of cellular compartments

In the interaction data gathered thus far, we wanted to investigate the presence of protein complexes, which may be connected to RTK signaling in the cell. To this end, we performed enrichment analysis of CORUM (Giurgiu et al., 2019) complexes for each RTK and then grouped the results based on the gene ontology cellular component (GOCC) annotations, if available in CORUM. Although many of the complexes had no localization annotations available, very thorough coverage of the cell was seen in the complexes that were able to be assigned to a locale (Fig. 2B). Curiously few strictly plasma membrane complexes were seen in the data. However, this may be in part due to imperfect coverage of GOCC annotations in CORUM and in part due to strict filtering applied to the data.

In total, 208 unique complexes were enriched in the data (Table S3), and we were able to assign probable localizations to 59 of these based on CORUM annotations. These assignments included five plasma membrane and eight ER complexes (two of which were specific ER-membrane complexes), five chromosomal complexes, and twenty-one other nuclear complexes. Other complexes enriched in the RTK interactor sets were two kinase maturation complexes and five different TNF-alpha/NF-kappa B signaling complexes. The most commonly enriched complex was the LTC-PLC-gamma-1-p85-GRB2-SOS signaling complex, which was enriched in 27 RTKs. The first of many ER protein complexes, coat protein complex II (COPII), was the second most common and was enriched with 21 RTKs. This complex shares many components with the two SEC23 complexes, which were also enriched in 21 RTKs.

Additionally, 26 nuclear complexes were identified. Based on existing knowledge and GO annotations, some of these complex components identified in this study do appear to shuttle between cytoplasm and nucleus, and even to the plasma membrane. However, the majority of the components in these complexes are strictly nuclear. Nuclear signaling is a well-documented, noncanonical mode of signaling for many RTKs (Carpenter, 2003; Krolewski, 2005; Massie and Mills, 2006; Schlessinger and Lemmon, 2006; Song et al., 2013). In our HCI data, we detected 93 strictly nuclear proteins with 40 different RTKs and 909 proteins with some activity in the nucleus according to GOCC classifications. Among strictly nuclear proteins, MER and FLT3 had the most interactions (22 interactions). In contrast, every RTK had interactors with some connection to the nucleus: DDR1, a collagen receptor, had the fewest (12, none of which were strictly nuclear). FGFR1 had the most (172), perhaps reflecting its known signaling functions in the nucleus (Myers et al., 2003; Stachowiak et al., 1996). While nuclear interactors can be explained by possible encounters during mitosis after nuclear breakdown, the data may also offer some additional context for possible connections between RTKs and nuclear signaling pathways.

The four identified HSP90-related complexes, which were significantly enriched with 47 different RTK baits, are of interest for the regulation of kinase activity. Considering the role of HSP90 in fostering and promoting proper protein folding and function, we next examined this link in detail. Of the 29 RTK baits that have previously been studied as potential interactors for the HSP90 complex (Taipale et al., 2012), 15 were strong interactors, 10 were weak interactors, and four were not interactors. Of the 3 HSP90 proteins of interest, CDC37, HSP90AA1, and HSP90AB1 were all identified with 11 RTKs, of which FGFR4 was not included in the Taipale et al study, and TYRO3 was classified as a weak interactor (Table S4). The nine others were strong interactors. CDC37 and HSPAA1 were identified as HCIs with LMR1. CDC37 alone was identified with all but 5 baits (Table S1). Therefore, our findings were consistent with those of Taipale et al. All three components were identified as HCIs for nine strong HSP90 interactors (Tables S1 and S4). These interactions were detected mainly via AP-MS, suggesting stable interactions. The only weak interactor that was detected with all three components, TYRO3, has since been linked to two HSP90 core interactor proteins (Li et al., 2018). FGFR4, which was not included in the Taipale et al. study, was identified with all three components by AP-MS, indicating that FGFR4 is a potential HSP90 interactor kinase.

### Enriched protein domains and functions of RTK interactors

Considering the enriched protein complexes identified, we next proceeded to investigate the domain composition of the individual HCI proteins (HCIPs). The top two domains identified by absolute counts were SH3 and SH2 (Fig. 3A). When considering only unique HCIPs, SH3, the protein kinase domain and the protein tyrosine kinase domain were the most common. All of these domains play prominent roles in kinase signaling (Mayer, 2001; Xin et al., 2013). The SH3 domain was identified 216 times in 39 unique HCIPs, whereas the SH2 domain was identified 180 times in 21 unique HCIPs.

**Figure 3.**
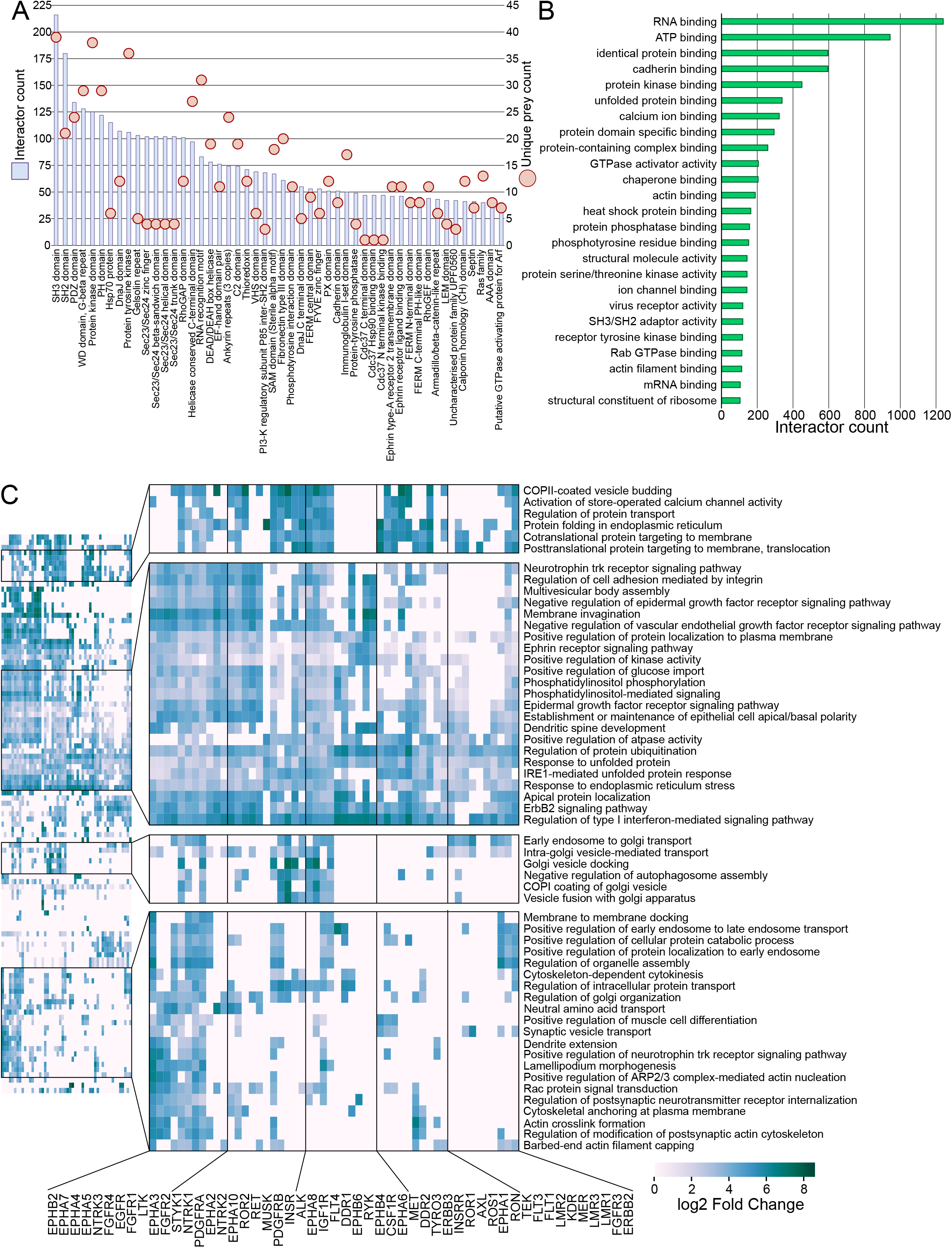
Characterization of RTK interactor proteins. A. Identified protein domains (PFam) of the RTK interactors. Blue bars (left Y-axis) denote the cumulative count of the corresponding domain, while light red circles (right Y-axis) denote count of unique prey proteins with the domain (i.e. SH3 domain was encountered 216 times in the data, but in 39 unique proteins, while SH2 domain was identified 180 in 21 unique HCIs). B. Significantly enriched (q<0.05) GO ‘molecular function’ annotations in the RTK interactors. C. Significantly enriched signaling pathways (Reactome) identified in each RTK interactome. Fold change values are shown in log2 scale.

Twenty-eight percent of all human proteins annotated with the protein tyrosine kinase domain were identified among the HCIPs, compared to 10% of proteins annotated with the protein kinase domain. SH2 domains suggest potential target proteins, since RTK activation via autophosphorylation induces the formation of SH2 domain binding sites (Lemmon and Schlessinger, 2010). Indeed, 43% of HCIPs with SH2 domains were previously known interactors of RTKs. To identify the specific functions these HCIPs participate in, we next examined GO molecular function terms associated with the identified HCIs. Similar to domains, the most common molecular functions associated with the HCIPs were related to protein kinase activities either directly (ATP binding), indirectly (protein kinase binding), or in a supporting role (heat shock protein binding) (Fig. 3B).

To investigate functional similarities and differences between RTKs based on their interactions, we next performed a GO biological process (BP) analysis and highlighted the most enriched (log2-fold change > 5) terms (Fig. 3C). We identified four clear groups of terms containing processes related to RTK functions. These included terms enriched in most RTKs, such as multiple signaling pathways, and groups of more specialized terms, such as processes related to vesicle trafficking between the Golgi apparatus and the endosomal system.

Many of these processes are interlinked with known RTK functions. The ERBB2 signaling pathway, for example, was significantly enriched in almost all RTKs. Similarly, the type I interferon signaling pathway was seen in all but three RTKs. As a further example, the Ephrin receptor pathway also contains the majority of RTKs. Given that among the pathways enriched with the highest fold change values, few are limited to individual receptors, the functional enrichment results further indicate that RTKs share many pathways through which signaling may occur depending on cellular conditions, possibly including crosstalk between the receptors.

We next examined how the enriched GOBP terms were represented among all previously known RTK interactors (Fig. S3A). In the analysis, some of the most common GOBP terms detected in our results, such as Signal transduction, protein phosphorylation and various signaling pathways (Fig. S3A, upper panel), were prominently featured in the database of known RTK interactors as well (Fig. S3A, lower panel). However, missing from the known interactors for many receptors were proteins connected to COPII vesicle coating and cargo loading, as well as PI3K activity regulation, all of which were common functions among the identified HCIs, possibly illustrating a gap in the previous knowledge in regard to such interactors. For example, COPII vesicle coating, budding, and cargo loading related proteins are missing from the known interactors of both RET and PDGFRB, but are found in our dataset in both pervanadate- and ligand-treated samples (Figure S3A, Table S1C).

### RTK interactors form protein clusters with distinctive functions

Previously, protein copurification was investigated in large-scale interaction studies to identify possible interactions between HCIPs. Affinity purification experiments showed that two proteins that purify together may indicate an interaction between them, such as a protein complex (Buljan et al., 2020; Mehta and Trinkle-Mulcahy, 2016; Yu et al., 2009). Therefore, to understand how the RTK HCIs detected in our study might interact with one another, we performed a cross-correlation analysis of both AP-MS (Fig. 4A, upper) and BioID (Fig. 4A, lower) data. In total, 2020 unique protein pair associations were detected through the two approaches (Table S5). A total of 105 of these were previously known interactions, and 130 protein-protein pairs were in the same reactome pathways. The analysis of random networks showed that this network was highly enriched in both known protein interaction pairs (Fig. S3B, top) and proteins in the same reactome pathways (Fig. S3B, bottom).

**Figure 4.**
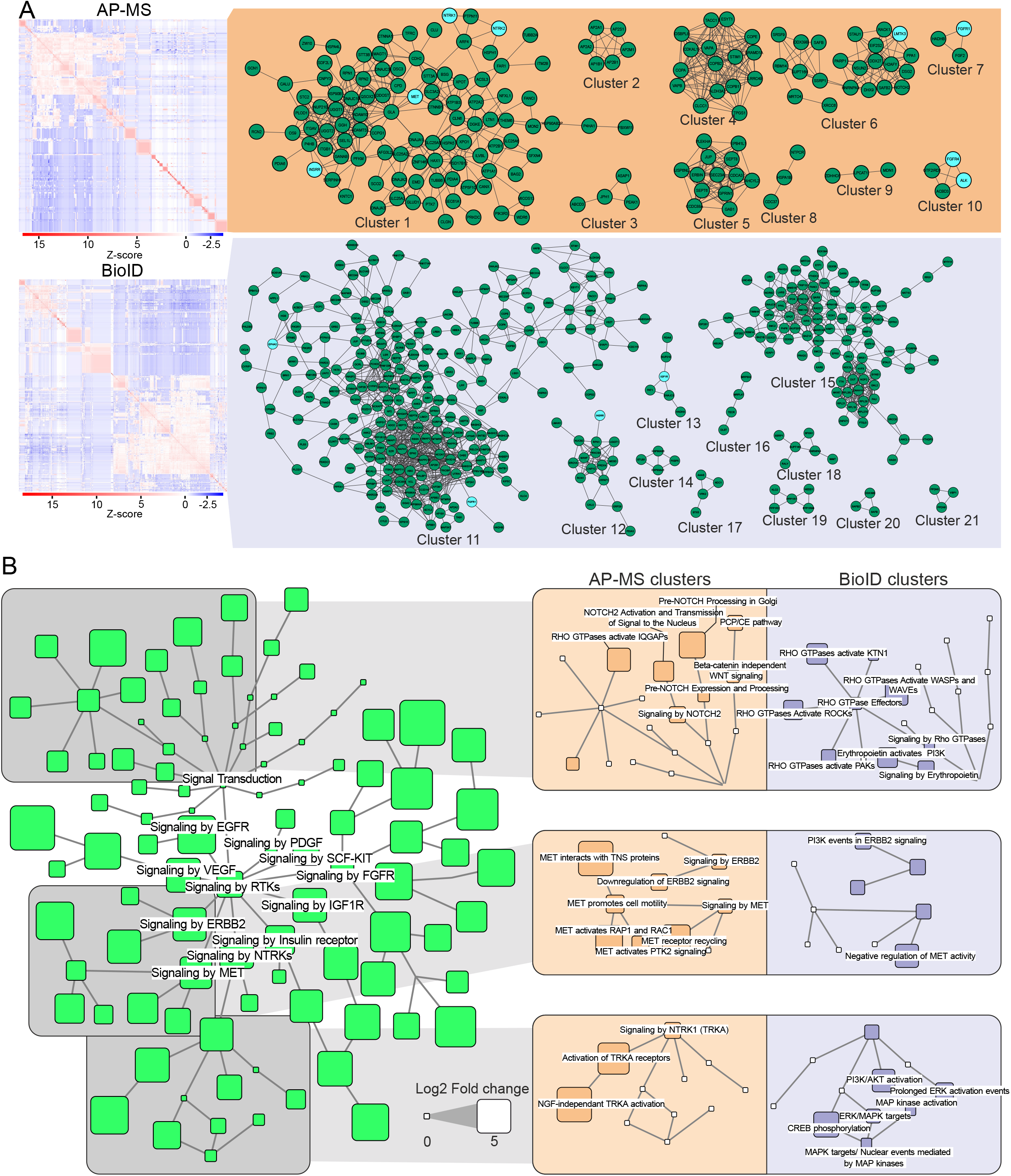
Functional clusters extracted from HCI Cross-correlation analysis. A. HCI-HCI association clusters identified via cross-correlation analysis of the identified RTK interactors. Clusters represent proteins, which are often co-purified in our experiments. Clusters were identified separately from the AP-MS or the BioID cross-correlation data. RTKs, if any, in the clusters are shaded light blue. B. Enriched (log2FC > 5, q<0.05) Reactome pathways in the identified association clusters. Nodes downstream from the signal transduction root node are shown. Node size corresponds to log2 fold change value of the pathway. Three pathway groups where AP-MS and BioID clusters had the most prominent differences in enrichment are further highlighted in the boxes with orange (AP-MS) and blue (BioID) background on the right.

From the dataset, 21 clusters with 3 or more proteins were identified (Fig. 4A). Of these, 10 were detected in AP-MS data and 11 in BioID. In total, 7 of the clusters featured one or more RTKs as well. Reactome pathway enrichment analysis was performed for each protein cluster to identify what functions each could participate in. The proteins of largest cluster detected in AP-MS data, cluster 1 (Fig. S3C, top left), functioned mainly in pathways such as small molecule transport, protein phosphorylation, and platelet signaling. The largest BioID cluster (11, Fig. S3C, bottom right) featured proteins in particular from multiple signaling pathways, as well as vesicle trafficking and endocytosis in particular (Table S6). From the clusters, also identified CORUM protein complexes (Fig. S3D). We filtered out all complexes from which less than 60 % of the components were identified, and removed overlapping complexes, keeping the more complete ones. This resulted in 30 protein complexes identified from the cross-correlation network.

We next linked the significantly enriched reactome pathway terms to the reactome hierarchy and extracted pathways linked to signal transduction (Fig. 4B). Several signaling pathways were enriched, particularly with AP-MS or BioID clusters. For example, RHO GTPase effector-related pathways were enriched in BioID clusters, while Notch and WNT signaling were enriched in AP-MS clusters. In the RTK pathways, we observed clear differences, particularly in the MET, ERBB2, and NTRK1 signaling pathway groups. These results suggest proximal RTK associations with functional protein networks related to RHO GTPase signaling, as well as MAPK and PI3K/AKT signaling. In contrast, the pathways enriched in the AP-MS clusters may indicate a more direct role for RTKs in protein clusters related to Notch and WNT signaling. The presence of core RTK pathways, such as TRKA receptor activation or MET signaling in the AP-MS clusters, strengthens the idea that RTKs have a more direct role in the pathways detected in AP-MS clusters.

Ephrin receptors A5, A6, A7, and A8 are some of the less well-studied RTKs (Fig. 1B). We therefore analyzed their interactomes and the interplay between these receptors. To focus on the common HCIs, we removed interactors seen with only one of these receptors (Fig. S4A). We identified the largest group of shared HCIs between EphA5 and EphA7, and there were 46 shared HCIs. In this group, we identified many other kinases, such as MAP4K and EphB4, and phosphatases, such as PTPN11 and PTPN13. We also identified nine HCIs shared between all four of the Ephrin receptors and 16 shared between EphA4, A7, and A8. The shared groups included multiple proteins that are integral to the function of RTKs, such as SEC23B, SEC24A, and SEC24B, which participate in coat protein complex II, which may indicate the use of COPII-coated vesicles in some portion of RTK membrane trafficking. When analyzing the interactions of enriched reactome pathways (Fig. S4B, left side), we indeed observed multiple transport pathways, including endosome-to-Golgi and Golgi-to-ER pathways. The interaction data therefore indicate possible RTK paths through the cell. When examining the enriched CORUM complexes in detail (Fig. S4B, right side), we identified the Wave2 complex and other actin dynamics-related factors, as well as oligosaccharyltransferase complexes responsible for co and posttranslational glycosylation of proteins in the ER lumen. Thus, the interactomics data may be used to identify core RTK interactors shared between subgroups of receptors and possible avenues for cooperative RTK actions.

### Potential substrates define RTK kinase activity

A heavy-labeled ^18^O-ATP-based *in vitro* kinase assay combined with LC-MS/MS (IVK, Fig. S5A) was used to characterize potential direct substrates of RTKs (Müller et al., 2016; Zhou et al., 2013). It is important to note that the kinases used in this method have access to not just their physiological molecular context but also proteins they may not normally encounter. Forty-five recombinant RTKs were used for experiments that included all RTK subfamilies. Any sites with a localization probability of under 0.75 were filtered out, as were sites seen in any of the control experiments, where recombinant kinase was not added. This resulted in a total of 2254 unique phosphorylated tyrosine sites, resulting in 7758 unique kinase-substrate interactions, or 10194 kinase-substrate phosphosrylation site relations (Fig. 5A, Table S7). Of the 10194, 6639 were novel, and 3555 were identified in a prior publication (Sugiyama et al., 2019), phosphoSitePlus, or phosphoELM. The number of identified sites varied widely between individual kinases, from nearly a thousand phosphotyrosine sites (982 substrate sites for EphB1) to fewer than five sites (Fig. 5C). A total of 1027 sites were detected with only one kinase, while others had up to 37 kinases (Fig. S5B). In contrast, in the control experiments without added kinase, a maximum of 5 phosphotyrosine sites were identified (Fig. S5B inset). Based on the PhosphoSitePlus database (Hornbeck et al., 2015), 1478 of the identified phosphorylation sites were previously reported, and the kinase responsible for phosphorylation was known for 124 of these sites. In 30 cases, we observed exactly the same kinase-substrate site interaction as was reported in PhosphoSitePlus (Table S7).

**Figure 5.**
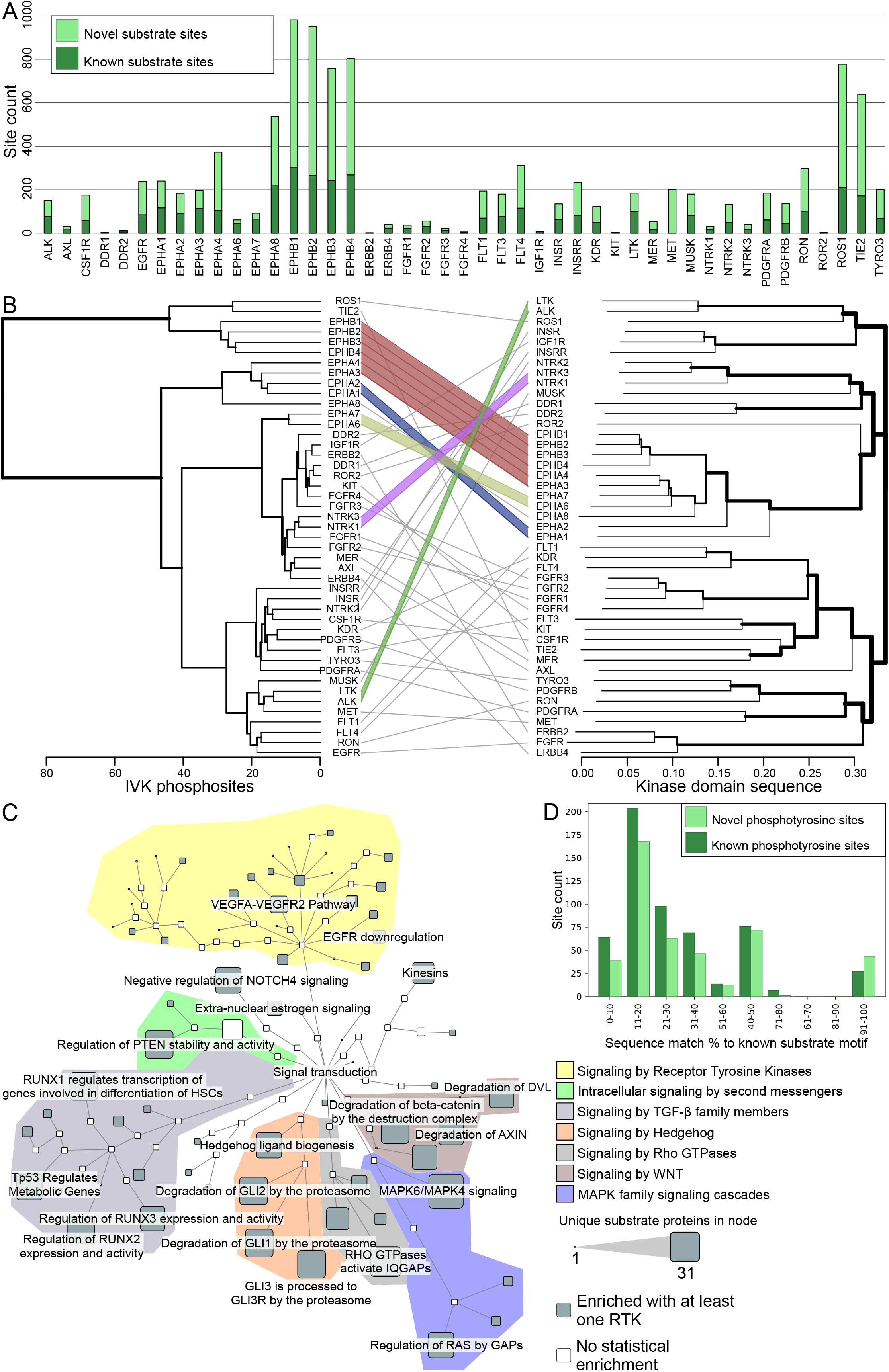
Characterization of RTK-specific phosphotyrosine sites. A. Phosphotyrosine sites identified in the IVK assay after filtering. Deeper shade of green corresponds to previously identified kinase-substrate relationships. B. Dendrograms of RTK clustering based on phosphosite identifications (left) compared to Clustal Omega clustering based on protein kinase domain sequence of the same RTKs (right). Colored lines denote baits in the same order in both clustering approaches. C. Statistically enriched Reactome terms in the identified RTK substrates. Size of the node corresponds to the number of unique substrates in the node, and nodes without significant enrichment are shaded white. Only subnodes of the signal transduction root node are shown. Colored areas denote different signaling pathway trees. D. Substrate site amino acid sequence compared to known phosphorylation motifs from human protein reference database (Peri et al., 2003). Data presented represents only the receptors, for which motifs were available in the database.

We performed clustering analysis of the detected phosphorylation sites to obtain an overall view of the RTK substrate profile, and the result was compared to the kinase domain sequence alignment tree produced by Clustal omega (Madeira et al., 2019) (Fig. 5B). Several kinase groups, the Ephrin receptor subfamily in particular, clustered together based on phosphosites, and most were close to their position in the kinase domain sequence-based tree. The main difference between the two dendrograms was the Ephrin receptor subfamily in the IVK analysis, which was divided in two: one group of four receptors and one of five receptors. Substrate site-based clustering indicated a distinction between the EphB1-4 group and EphA1-A8 group, while the subfamily according to the kinase domain sequence is in one well-defined branch. The IVK analysis results were also compared with clustering results from the AP-MS and BioID data (Fig. S5C), and no strict similarity in the interactor profiles of receptors in the Ephrin subfamily was observed. This may be due to two factors. First, the number of identified phosphosites or HCIs per RTK varies, and when a few are identified, the clustering algorithm does not work. Second, substrates may also vary significantly within receptor families. However, when all three approaches (AP-MS, BioID, and IVK) produced similarly unorganized clusters, it seems plausible that RTK substrate and interactor profiles may vary as much within subfamilies as between them. On the other hand, similarities detected between RTK substrates suggest a similarity among some functions. One such case is KDR and PDGFRB, and similarities in their IVK substrate profiles may indicate functional similarities. Indeed, the two proteins share 90 previously known interactors (Table S1) and 53 phosphosites detected in our IVK experiments, indicating a strong basis for overlapping functions.

A reactome enrichment analysis was used to link the identified RTK substrate proteins to functional networks. We focused on pathways linked to signal transduction to study the possible significance of the kinase-substrate relationships in cellular signaling networks (Fig. 5C, Table S8). While signaling by RTKs was very prominent, the signaling pathways with the highest number of identified proteins were ‘MAPK6/MAPK4 signaling’ (31 substrate proteins) and ‘RHO GTPases Activate Formins’ (27 proteins). The most commonly enriched pathway was the ‘VEGFA-VEGFR2 pathway’, which was seen with 38 of the 45 kinases used, but there were only 15 unique substrate proteins. In particular, the enrichment of the Wnt, TGF-β and MAPK signaling pathways may be due to a previously known link between RTKs and regulation of these three signaling pathways (Billiard et al., 2005; Heldin and Moustakas, 2016; Katz et al., 2007; Krejci et al., 2012; Shi and Chen, 2017). When examining the identified substrates in detail, out of the seven pathway groups emphasized in Fig. 5C, TGF-β had the highest number of substrates (Fig. S5D). Our data may therefore provide further information of these links.

Of the 10194 RTK-substrate site relationships identified, 3566 were found in one or more of the three databases used to identify known phosphorylation sites of these kinases (PhosphoSitePlus, phosphoELM, Sugiyama et al., 2019. A further 5 sites had identical surrounding +/-7 amino acids as in a previously identified substrate site. To further query whether the novel sites shared similarity with the previously identified, we next compared the known and novel substrate sites to known phosphorylation motifs from the human reference protein database. The motif match percentage profile between known and novel phosphorylation sites is generally of the same shape, however we identified more perfect matches to the annotated motifs in the set of novel substrate sites (Fig 5D).

### Kinase activity-deficient mutants reveal activity-dependent functions

KD RTK mutants were used to understand which interactions might be dependent on RTK protein kinase activity. We performed AP-MS and BioID experiments with KD mutants and compared the results to the WT RTK results. The kinase domain in the mutants was deactivated with a point mutation that introduced bulk into the ATP binding pocket. The number of HCIs we identified varied widely depending on the receptor (Fig. 6A, Table S1). Some WT RTKs, such as AXL, EphA7, and MER, had more HCIs than their KD counterparts, whereas in others, DDR2 in particular, the KD mutant had more HCIs.

**Figure 6.**
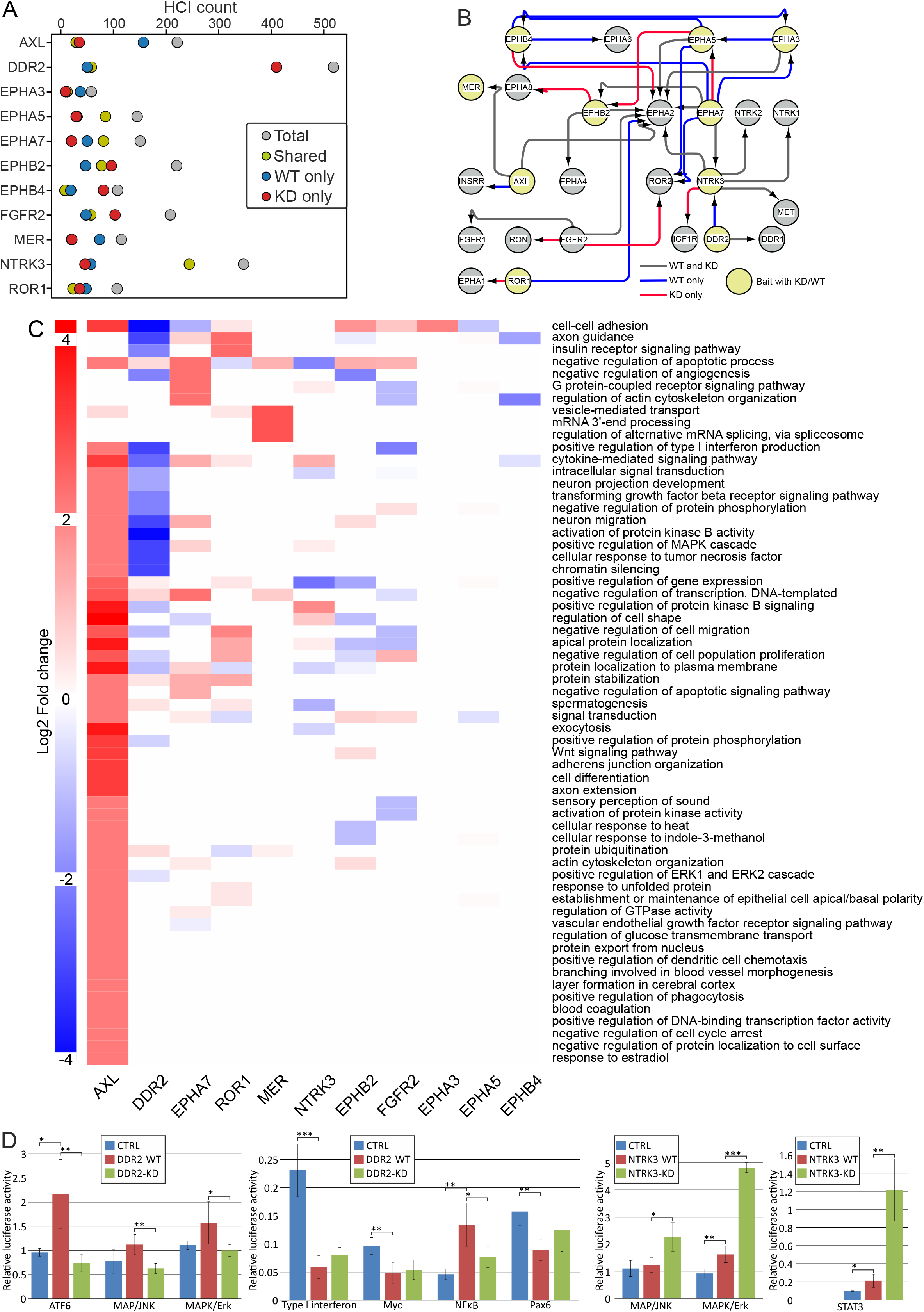
Assessment of differences in wild type and kinase dead RTK mutants. A. HCI counts per WT / KD pair. Total HCI number is shown in grey, while number of shared HCI proteins is in yellow, WT only HCIs in blue, and KD only HCIs in red. B. Bait-bait interactions of the WT / KD baits. Shown are all RTKs found in WT/KD HCI data, but interactions are shown only for those with WT and KD constructs. Grey arrows depict preserved interactions, while blue ones are interactions that are lost in KD data, and red denotes interactions only seen in KD data. C. GO biological process change in KD data. Values are log2 fold change in KD compared to WT, where positive values reflect higher representation in KD data. D. Comparison of effects of DDR2 and NTRK3 WT and KD on activity of cellular signaling pathways. *:p<0.05, **:p<0.01,***:p<0.001

Considering the prominent role of RTK-RTK interactions in the WT data (Fig. 2A), we first identified whether these interactions were gained or lost with the KD mutant (Fig. 6B). While many interactions were lost, a similar number was also gained, suggesting that the ability of KD mutants to associate with other RTKs in general is not significantly impeded by the inability to bind ATP. However, individual RTKs such as EphA3, A5, A7, and EphB4 seem to lose many interactions with other members of the Eph subfamily. Three of these interactions were detected only by AP-MS, two only by BioID, and two by both methods. This finding may indicate a reduced capacity of these RTKs to form heterodimers.

We then decided to sum up the lost or gained interactions by characterizing them via GOBP terms (Fig. 6C). To isolate pathways that may be lost or gained by the KD mutants, we calculated fold change values for the KD experiments using WT experiments as background. The results determined which terms were proportionally better represented in KD mutant HCIs (such as cell-cell adhesion in AXL KD) and in WT HCIs (such as cell-cell adhesion in DDR2). These results show that although the WT AXL has more HCIs than the KD counterpart, the different proteins do not concentrate heavily on any specific GOBP annotation; hence, fewer GOBP terms are overrepresented in the WT data than in the KD HCI set.

Likewise, although the DDR2 KD mutant had a much higher number of interactors than the WT counterpart, very few pathways had a positive fold change. DDR2 is a part of the DDR subfamily of collagen receptors. The loss of cell-cell adhesion pathways in the KD mutant (Fig. 6C) therefore suggests the loss of this core function. This finding together with fewer enriched pathways in general and the exceptional number of HCIs identified in BioID experiments for the DDR2 KD mutant (Table S1) indicates a proximity to a wider variety of proteins, possibly stemming from irregular cellular localization for the KD mutant.

To identify if the KD mutation had an identifiable effect on a transcription level, we next performed a luciferase assay panel measuring transcription factor activity as a response to the transfected kinase (Fig. 6D). With DDR2, where we saw the largest difference between WT and KD interactomes, we also detected significant changes in transcription factor activity. ATF6, MAP/JNK, MAPK/Erk, and NFKB pathways showed a significantly different response between the KD mutant and the wild type kinase. In all cases, the response of the KD-transfected cells was lower than that of WT. In contrast, with NTRK3 we saw significantly different responses in MAP/JNK, MAPK/Erk, and STAT3 pathways. However, in these cases, the wild type elicited a weaker response. Together, the data from the performed luciferase assay suggests that WT DDR2 and NTRK3 may produce opposing effects on MAP/JNK and MAPK/Erk signaling pathways.

### Known roles of EGFR identified via interactome analysis

After assessing the data produced in this study as a whole, the interactomes of singular receptors was focused on. To validate our results, we first focused on the well-known receptor EGFR (Fig. S6). Among the EGFR HCIs, we identified 94 previously known interactors, including other kinases (e.g., EphA2 and ERBB4) and phosphatases, such as PTPN1 and PTPN11. In addition to known interactors, we identified 137 novel interactors (Fig. S6A). GOBP enrichment analysis was used to discover which processes were driven by known and novel interactors. In this set of enriched GOBP terms, the most commonly identified ones were often driven by a mixture of known and novel interactions (Fig. S6B). To see how the novel interactors relate to the known ones, we next identified the previously known interactions between the known and novel HCIPs (Fig. S6C). From these data, we could see that the novel interactors often act as bridges or network hubs between different known interactors, such as MAP3K7, LTN1, or XPO1. Furthermore, some of the novel interactors are closely related to the known ones. For example, although interaction with ABI1 is included in the combined database of previously known interactions, ABI2 was not. Similarly, VAPA is in the known interaction database, whereas VAPB is not. To validate interactions identified by our approach, we chose 9 AP-MS-detected HCIPs at random for CO-IP analysis. Of these, only one failed to show a clear interaction in the resulting blot (Fig. S6D, left). As two of the proteins chosen were also detected in NTRK3 AP-MS data (SEL1L and SEC61A), we chose to further ensure the reliability of the method by performing a CO-IP experiment targeting these two as well. (Fig. S6D, right).

In the enriched biological processes (Fig. S6B), we identified terms driven only by known interactors, such as clathrin-dependent endocytosis, terms driven by both, such as the VEGFR signaling pathway, and functions related to novel interaction partners, such as the positive regulation of arp2/3 complex-mediated actin nucleation. Clathrin-mediated endocytosis of EGFR is a major active pathway of receptor internalization (Sigismund et al., 2008). After endocytosis, EGFR may be either recycled back to the membrane or degraded, depending on ubiquitinylation. In addition to the enriched clathrin-dependent endocytosis identified by GOBP analysis of the EGFR interactome, we also detected multiple ubiquitinylation proteins. Six of these (CTNNB1, OS4, PRKDC, UBE2M, UBE2N, and SH3RF1) were previously documented EGFR interactors, while another four (CAND2, CDCA3, LTN1, and TRIM13) were novel interactors. Our data therefore provide additional support for the previously known interactors and molecular processes of EGFR. Furthermore, the interactome provides an additional molecular context for EGFR actions and dynamics with possible connections to novel functions.

### Characterization of the novel EphA7 interactome and phosphorylome

EphA7 is one of the least well-characterized members of the Ephrin receptor subfamily, with only 12 known interactors in IntAct. We therefore more closely analyzed the identified interactions and substrates of EphA7. WT EphA7 was analyzed together with the KD mutant to gain insights into the functions of WT EphA7 and how these functions are impacted by the loss of kinase activity (Fig. 7A). We divided the interactor proteins into the following groups: WT only, KD only, and shared proteins. In total, we identified 131 HCIs for the WT protein and 101 for the KD mutant. Of the 12 previously known interactors, we detected 3 in our experiments: EphA3 was only in WT, EphA2 was in WT and KD, and GNB1 was in KD only. Although EphA2 was detected in both, in the KD experiments, it was only seen by BioID, perhaps indicating loss of heterotypic complex formation with EphA2. The formation of heterotypic complexes is a well-documented behavior of the Eph subfamily of receptors (Janes et al., 2011), and given the detection of EphA5 in KD AP-MS data only, it seems unlikely that the ability to form these complexes is completely destroyed by the KD mutation.

**Figure 7.**
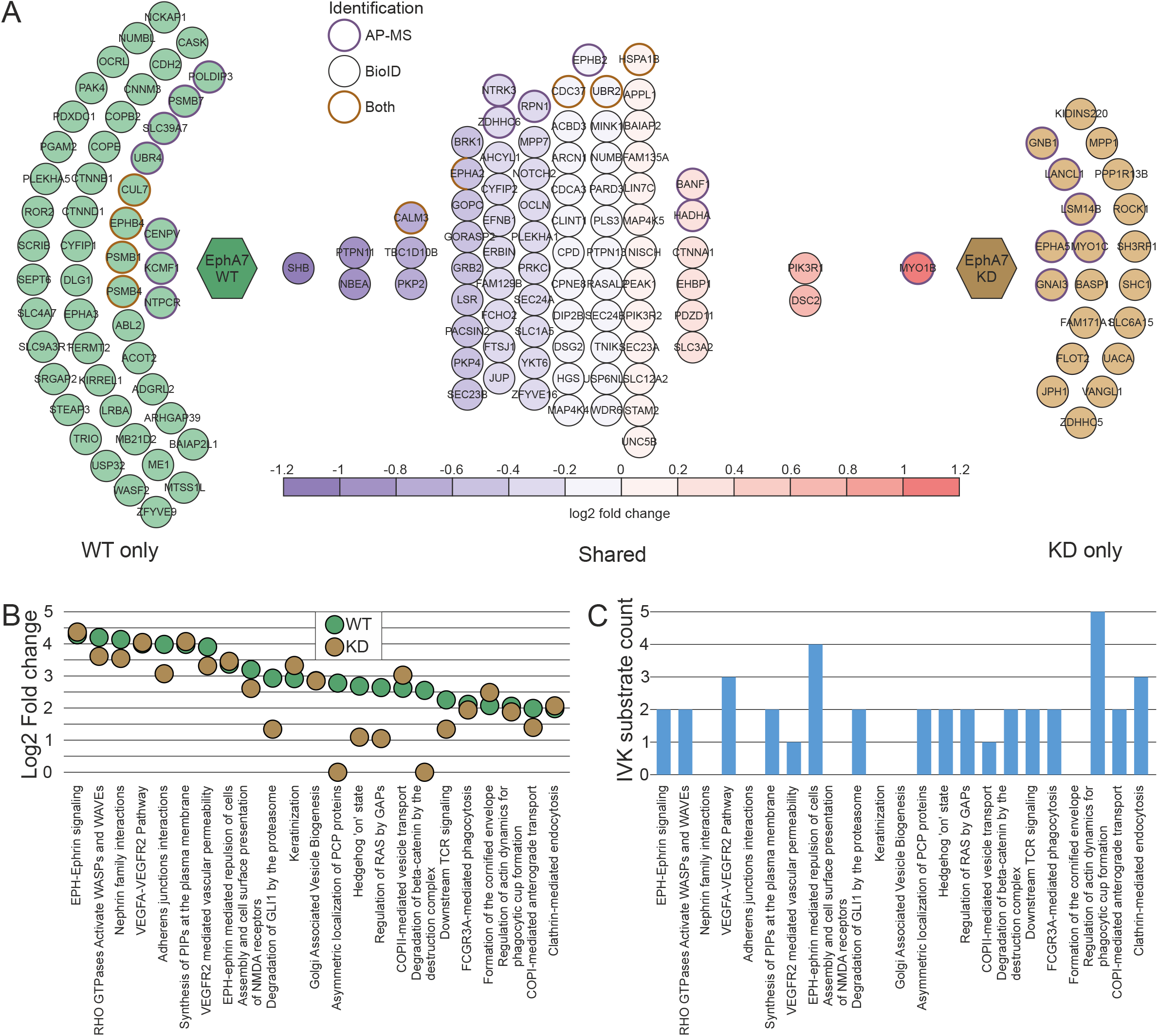
EphA7 interactome and phosphorylome analysis. A. EphA7 WT (left) and KD (right) HCIs. Shared HCIs are in the middle arranged according to log2 fold change values. HCIs identified in AP-MS are marked with a violet rim, BioID with black rim, and orange rim marks HCIs detected with both approaches. For the shared interactors, a bait-normalized fold change value was calculated. Four HCIs, CALM3, CDC37, UBR2, and HSPA1B were identified in both WT and KD experiments with both AP-MS and BioID methods. For these, the fold change values in the different experimental approaches were within 0.1 of each other, and thus the value used was an average of both. EphA2 was detected via AP-MS and BioID with WT EphA7, and with only BioID with KD EphA7. B. Significantly enriched (q<0.01) Reactome pathways in EphA7 WT data. Log2 fold change values are shown for both WT (green) and KD (orange). The KD values used did not undergo filtering to avoid eliminating smaller effects. C. Counts of substrates identified with the IVK method in the Reactome pathways enriched in EphA7 WT HCI data.

In the shared group, three proteins (SHB, PTPN11, and NBEA) clearly associated more with WT EphA7 than with the KD mutant, and one (MYOB1) associated more with the mutant. PTPN11 is a phosphatase with known roles in EphA2 and WNT signaling (Miao et al., 2000; Noda et al., 2016). This, together with EphA2 detection in the WT AP-MS data, also indicates potential cooperation by these two RTKs and the loss of this function when the activity of the kinase domain is compromised. Multiple proteasomal components (PSMB1, 4, and 7) and ubiquitinylation proteins (CUL7, KCMF1, and UBR4) were only detected in WT experiments. Their presence may mean that proteasomal degradation of EphA7 is the endpoint of the receptor, as it is for some other RTKs (Geetha and Wooten, 2008; Jeffers et al., 1997). Moreover, the absence of these proteins in the KD data may indicate that the process is dependent on RTK kinase activity.

We next determined how the differences in HCIs affected the most enriched reactome pathways in the EphA7 data. The Ephrin signaling and VEGFA-VEGFR2 pathways were represented by nearly identical proportions of HCIs in both the KD and WT experiments. However, differences could be seen in other pathways, especially in planar cell polarity (PCP) protein localization and various signaling events. It is possible that the KD mutation does not affect the association with proteins related to many of the signaling pathways but does affect the association with specific participants in the signaling cascades, such as the aforementioned SHB and PTPN1.

To understand how EphA7 affects the pathways it is most strongly linked to in our AP-MS and BioID data, we combined the data with substrates identified by the IVK method, and found EphA7 substrates in most of the pathways were enriched in the HCI data. Of the pathways that differed most between the WT and KD experiments, degradation of beta-catenin by the destruction complex, degradation of GLI by proteasomes, asymmetric localization of PCP proteins, Hedgehog ‘on’ state, and the regulation of RAS by GAPs all had identified phosphosites in the IVK data (Fig. 7B-C).

Taken together, data produced by our systematic approach to identify interactors and phosphorylation targets of EphA7 suggest that the KD mutation does not hinder the association with proteins in Ephrin signaling pathways but may affect specific receptor localization, as reflected by the reduced number of proteins identified in other signaling pathways. The IVK data can additionally be used to identify specific target candidates for EphA7 in the Ephrin and VEGFA signaling pathways. Furthermore, the HCIs identified for WT EphA7 suggest that proteasomal degradation may be the termination point receptor signaling, and their absence in the KD data suggests that the process is dependent upon the kinase activity of EphA7.

## Discussion

Here, we present the comprehensive interactome and phosphorylome of human RTKs. RTKs play key roles in initiating a complex web of signaling cascades. While many have been well studied (Fig. S1B), detailed and systematic knowledge of the roles and actions of a large proportion of RTKs, such as many Ephrin receptors, is lacking. In this study, we used three complementary approaches to understand RTK functions: AP-MS to capture stable interactions and complex stoichiometries, BioID to capture transient interactions and molecular context, and IVK to identify RTK substrates. To date, this dataset is the most comprehensive resource of RTK interactions and substrates. The data introduced here provide information about protein complexes (AP-MS), the surrounding molecular landscape (BioID), and signaling activity (IVK). Overall, these three approaches can be used to characterize and introduce additional context for well-known receptors (Fig. S6), discover the functions of less well-known receptors (Fig. 7), and identify possible active roles for RTKs in signaling networks via substrate information (Fig. 5). The data supplement the scarce information available for some RTKs, and for the whole kinase family, these data underscore the interactions within and across subfamilies. While the interconnectedness of RTK signaling networks is a well-known feature of these receptors (Kholodenko et al., 2010; Paul and Hristova, 2019), the data presented in this study supply additional molecular context for the signaling networks and indicate probable avenues of information flow. The interactomics insights gained here highlight the role of RTKs as important intersections in an increasingly complicated landscape of cellular signaling networks.

Despite the comprehensiveness of the results presented here, our model does have several limitations inherent to large-scale high-throughput proteomic studies. The results do not capture all context-dependent interactions. Our use of pervanadate to ensure the capture of active-state interactions does alter the specific molecular landscape of the cells, and thus, the detected interactions do not necessarily reflect in vivo activation of RTKs. Furthermore, the isoforms expressed in various cell populations may differ from the isoforms used here, and N-terminal tagging of these constructs may affect some interactions or protein stability. Indeed, not all RTKs are physiologically expressed in HEK293 cells (Table S1B), nor does this cell line represent all common cell types. Therefore, the new roles indicated by interaction data presented in this study, while suitable for hypothesis generation, should not be taken to confirm any novel functions of RTKs. The IVK method has the caveat of using recombinant RTKs and giving each kinase access to more than their physiological molecular context. Although nuclear proteins are not specifically solubilized, substrates available in this method may cover e.g. membrane domains or structures from which RTKs are normally excluded from.

In summary, the study describes the RTK molecular context and interactomics landscape, as seen from the perspective of AP-MS and BioID methodology, and the phosphorylome as identified by *in vitro* kinase assays. The combined knowledge of the multifaceted dataset presented may best be used as a potential pool for each RTK and be combined with additional application-specific information, such as data on specific cancer types or drug applications, to generate testable hypotheses of molecular systems surrounding RTKs. The data may also be used to gain insight and context into known functions of well-studied kinases, such as EGFR (Fig. S6), or to derive indications of possible roles for less well-known RTKs, such as EphA7 (Fig. 7). Furthermore, systemic insights can be gained by studying the connections within groups of receptors, of which we chose EphA5-A8 as an example subgroup (Fig. S4). The knowledge presented herein emphasizes common functions between RTKs and the landscape that they share with other signaling pathways. The three perspectives of the data presented here, stable interactions (AP-MS), proximal and transient interactions (BioID), and kinase-substrate relationships (IVK), together form a comprehensive molecular environment that can serve as a foundation for a systemic view of RTK signaling pathways and networks.

## Materials and methods

### RTK constructs

RTK constructs were obtained from three sources: 7 were gifts from William Hahn & David Root (Addgene plasmid # 23914, 23906, 23900, 23910, 23892, 23883, 23891) (Johannessen et al., 2010). 15 were from ORFeome collection (ORFeome and MGC Libraries; Genome Biology Unit supported by HiLIFE and the Faculty of Medicine, University of Helsinki, and Biocenter Finland), and 29 from a collection published previously (Varjosalo et al., 2013a), and 2 were synthesized by Genscript. RTKs were cloned into MAC-TAG-C expression vector (Liu et al., 2018) and pcDNA™-DEST40 (Thermo Fisher Scientific) via gateway cloning.

### Cell culture

Stable cell lines inducibly expressing the MAC-tagged RTK baits, Flp-In 293 T-REx cell lines (Cultured in DMEM (4.5 g/l glucose, 2mM L-glutamine) supplemented with 50 mg/ml penicillin, 50 mg/ml streptomycin and 10 % FBS) were co-transfected with the expression RTK vector, and pOG44 vector (Invitrogen) using FuGENE 6 transfection reagent. Cells were selected with 50 ml/ml streptomycin and 100 μg/ml hygromycin for two weeks, starting two days after transfection. Positive clones were pooled and amplified in 150 mm plates. Each cell line was expanded to 80 % confluence in 20 × 150 mm plates. Ten of the plates were used for AP-MS approach and 10 for BioID. For AP-MS, expression of the bait protein was induced with 1 μg/ml tetracycline 24 h prior to harvesting. With BioID, 50 μM biotin was added to the plates in addition at induction with tetracycline. Pervanadate treatment was performed at a concentration of 100 µM for 15 minutes prior to harvesting. Cells from 5 × 150 mm fully confluent plates (∼ 5 × 10^7^ cells) were harvested on ice and pelleted as one biological sample, thus each bait protein had two biological replicates for each of the two approaches. Samples were snap frozen and stored at -80°C. Tetracycline concentration of 1 μg/ml was used to produce expression levels corresponding to expression levels similar to endogenous (Glatter et al., 2009; Varjosalo et al., 2013a; Varjosalo et al., 2013b; Yadav et al., 2017).

For ligand experiments, the cells were treated with the ligand of the expressed RTK (EGF, 10 ng/ml; FGF, 10 ng/ml; IGF 20 ng/ml; HGF 50 ng/ml; NT-3 10 ng/ml; PDGF-BB 10 ng/ml; GDNF 10 ng/ml; All from R&D systems). Ligand treatment was started at the time of tetracycline induction, 24 h before harvesting. Ligand experiments were carried out for 8 RTKs (EGFR, FGFR1, FGFR4, IGF1R, INSR, INSRR, PDGFRB, and RET).

### Affinity purification of RTK interactors

For AP-MS, samples were lysed in 3 ml of ice-cold lysis buffer 1 (0.5 % IGEPAL, 50 mM Hepes (pH 8.0), 150 mM NaCl, 50 mM NaF, 1.5 mM NaVo_3_, 5 mM EDTA, with 0.5 mM PMSF and protease inhibitors (Sigma-Aldrich)).

For BioID approach, cell pellets were thawed in 3 ml of ice-cold lysis buffer 2 (0.5 % IGEPAL, 50 mM Hepes (pH 8.0), 150 mM NaCl, 50 mM NaF, 1.5 mM NaVo_3_, 5 mM EDTA, 0.1 % SDS, with 0.5 mM PMSF and protease inhibitors (Signa-Aldrich). Lysates were sonicated and treated with benzonase (Bio-Rad).

Lysates were centrifuged at 16000 × G for 15 minutes, after which the supernatant was centrifuged for an additional 10 minutes to obtain cleared lysate. This was then loaded consecutively on spin columns (Bio-Rad) containing 200 μl Step-Tactin beads (IBA,GmbH) prewashed with lysis buffer 1. The beads were washed thrice with 1 ml of lysis buffer 1, and 4 × 1 ml of wash buffer (50 mM Tris-HCl, pH 8.0, 150 mM NaCl, 50 mM NaF, 5 mM EDTA). After the final wash, beads were resuspended in 2 × 300 μl elution buffer (50 mM Tris-HCl, pH 8.0, 150 mM NaCl, 50 mM NaF, 5 mM EDTA, 0.5 mM biotin) for 5 minutes, and eluates were collected into 2 ml tubes. Cysteine bonds were then reduced with 5 mM Tris(2-carboxyethyl) phosphine (TCEP) for 30 minutes at 37 °C, followed by alkylation with 10 mM iodoacetamide for 20 minutes in the dark. Proteins were then digested to peptides with sequencing grade modified trypsin (Promega V5113), at 37 °C overnight.

The following day quenching was done with 10 % TFA, and the samples were desalted with C18 reversed-phase spin columns. These columns were first washed 3 × 100 ul of 100 % acetonitrile (ACN), and equilibrated with 3×100 ul of buffer A(0.1%TFA, 1% ACN). This was followed by 4 × 100 ul of wash buffer (0.1 % TFA, 5 % ACN). Peptide samples were then loaded 300 ul at a time, followed by 4 × 100 ul washes with wash buffer. Elution was done with 3 × 100 ul of elution buffer (0.1 % TFA, 50 % ACN). The eluted peptide sample was then dried in a vacuum centrifuge and reconstituted to a final volume of 30 μl in buffer A.

### In-vitro kinase assay

HEK 293 cells were cultivated in DMEM (GE Healthcare), supplemented with 10% foetal bovine serum (FBS) and antibiotics (penicillin, 50 ug/mL and streptomycin, 100 μg/mL). Cells on a plate were washed with PBS, dislodged with PBS and EDTA, and collected with centrifugation at 1,400 × g for 5 min before lysis. Cell lysate was prepared by lysing the pelleted cells with buffer containing 50 mM Tris-HCL, pH 7.5, 150 mM NaCl, 5 mM EDTA, 1% NP-40 (Invitrogen, Thermo Fisher Scientific), and protease inhibitors cocktail (Sigma) on ice. The cell debris was cleared by centrifugation at 16,000 × g for 10 min. The protein contents were measured using a BCA protein assay kit (Pierce, Thermo Scientific) and the cell fractions were stored at -80 °C.

Cell fractions were thawed on ice and endogenous kinases were inhibited with 5′-[p-(fluorosulfonyl)benzoyl]adenosine (FSBA; Sigma-Aldrich) in DMSO at a final concentration of 1 mM FSBA and 10% DMSO in Tris-HCL, pH 7.5 for 1 h at 30 °C. Excess FSBA reagent was removed by ultracentrifugation with 15 mL 10 K MWCO Amicon® Ultra-4 centrifugal filter units (Merck) at 3,500 × g at RT. Proteins were washed 4× the initial volume with kinase assay buffer (50 mM Tris-HCl, pH 7.5, 10 mM MgCl2, 1 mM DTT), adjusted to 2 mg/ml and stored on ice. For kinase reaction, 200 μg (100 μl) of FSBA-treated cell lysate was incubated with 1 μg of kinase (Life Technologies) and 1 mM γ[^18^O_4_]-ATP (Cambridge Isotope Laboratory) in 30 °C for 1 hour. For negative control experiments, 200 μg of FSBA-treated cell lysate was incubated with 1 mM γ[^18^O_4_]-ATP in the absence of added kinase. Reactions were halted with 100 μl of 8 M urea.

Prior to digestion, the proteins the samples were reduced with 5 mM Tris(2-carboxyethyl)phosphine (TCEP; Sigma-Aldrich) for 20 minutes in 37 °C, and then alkylated with 10 mM iodoacetamide (IAA; Sigma-Aldrich) for 20 min in room temperature in the dark. 600 μl of ammonium bicarbonate (AMBIC; Sigma-Aldrich) was added to dilute urea before trypsin digestion. Sequencing Grade Modified Trypsin (Promega) was then used to get a 1:100 enzyme:substrate ratio and the samples were incubated overnight at 37 °C. After digestion, the samples were desalted with C18 macrospin columns (Nest Group).

The macrospin columns were first conditioned by centrifuging 200 ul of 100% ACN through at 55 × g, followed by 200 ul of water. Column was then equilibrated twice with 200 ul of buffer A(0.1 % TFA, 1 % ACN). Sample was then added 100 ul at a time, and washed twice with 200 ul of buffer A. Finally, the sample was released with 3×200 ul of elution buffer (80 % ACN, 0.1 % TFA).

Phosphopeptide enrichment was performed using immobilized metal ion affinity chromatography with titanium (IV) ion (Ti4+-IMAC). The IMAC material was prepared by following the steps of the protocol published previously (Zhou et al., 2013). For enrichment of phosphopeptides, the Ti4+-IMAC beads were loaded onto GELoader tips (Thermo Fisher Scientific). The material was then conditioned with 50 μl of conditioning buffer (50 % CH_3_CN, 6 % TFA) by centrifuging at 150 g until all of the buffer had gone through. The protein digests were dissolved in a loading buffer (80 % CH_3_CN, 6 % trifluoroacetic acid (TFA)) and added into the spin tips and centrifuged at 150 g until all had gone through. The columns were then washed with 50 μl of wash buffer 1 (50 % CH_3_CN, 0.5 % TFA, 200 mM NaCl), followed by 50 μl of wash buffer 2 (50 % CH_3_CN, 0.1 % TFA), and finally the bound phosphopeptides were eluted with 10% ammonia, followed by a second elution with elution buffer (80 % CH_3_CN, 2 % FA). Samples were then dried in a vacuum centrifuge and reconstituted to a final volume of 15 μl in 0.1 % TFA and 1 % CH_3_CN.

### Liquid chromatography-Mass spectrometry (LC-MS)

The LC-MS/MS analysis was performed on Q-Exactive or Orbitrap Elite mass spectrometers using Xcalibur version 3.0.63 with an EASY-nLC 1000 system attached via electrospray ionization sprayer (Thermo Fisher Scientific). For each sample two biological replicates were used. Peptides were eluted and separated with C-18-packed precolumn and an analytical column, using a 60-minute buffer gradient from 5 to 35 % buffer B, followed by 5-minute gradient from 35 to 80 % buffer B, and a 10-minute gradient from 80 to 100 % buffer B at a flow rate of 300 nl/min (Buffer A: 0.1 % formic acid in 2 % acetonitrile and 98 % HPLC-grade water; buffer B: 0.1 % formic acid in 98 % acetonitrile and 2 % HPLC-grade water). Four microliters of peptide sample was loaded for each analysis from an enclosed, cooled autosampler. Data-dependent FTMS acquisition was in positive ion mode for 80 minutes, and a full scan from 200 to 2000 m/z with a resolution of 70 000 was performed, followed by top 10 CID-MS2 ion trap scans with a resolution of 17500. Dynamic exclusion was set to 30 s.

The Acquired MS2 spectral data files (Thermo RAW) were searched with Proteome Discoverer 1.4 (Thermo Scientific) using SEQUEST search engine against human protein database extracted from UniProtKB (https://uniprot.org) on 26.03.2019. For the searches, trypsin was set as the digestion enzyme with a maximum of two missed cleavages permitted. Precursor mass tolerance was set to +/-15 ppm, and fragment mass tolerance to 0.05 Da. Carbamidomethylation of cysteine was defined as a static modification, and oxidation of methionine, and biotinylation of lysine and N-termini as variable modifications.

For the kinase assays, LC-MS/MS analysis was performed as before, except peptide separation gradient was a 120-minute linear gradient. The IVK raw data files were processed with MaxQuant version 1.6.0.16 (Cox and Mann, 2008). MS spectra were searched against the human component of the UniProtKB database (release 2017_12 with 20192 entries) using the Andromeda search engine (Cox et al., 2011). Carbamidomethylation (+57.021 Da) of cysteine residues was used as static modification. Heavy phosphorylation of serine/threonine/tyrosine (+85.966 Da) and oxidation (+15.994 Da) of methionine were used as dynamic modifications. Precursor mass tolerance and fragment mass tolerance were set to less than 20 ppm and 0.1 Da, respectively. A maximum of two missed cleavages was allowed. The results were filtered to a maximum false discovery rate (FDR) of 0.05. Processed data was analyzed manually and filtered based on localization probability with a cut-off at 0.75. Any phosphotyrosine sites that were identified in control experiments without added kinase were also discarded.

### Data filtering steps

Significance Analysis of INTeractome (SAINT) express version 3.6.0 (Choi et al., 2011) and Contaminant Repository for Affinity Purification (CRAPome, http://www.crapome.org) (Mellacheruvu et al., 2013) were used as statistical tools for identification of specific high-confidence interactions from AP-MS and BioID data. 70 control runs with MAC-Tagged GFP were used as controls for SAINT analysis. Identifications with a SAINT-assigned bayesian FDR >= 0.05 were dropped, as well as any proteins that were detected in >= 20 % of CRAPome experiments, unless the spectral count fold change was over 3 when compared to CRAPome average. The remaining high-confidence interactors (HCIs) were then used for further analysis. For the IVK method, any phosphosite with <75 % localization probability as assigned by MaxQuant were discarded, as were sites that were detected in any control sample.

### Databases

Known interactors were mapped from BioGRID (only experimentally detected interactions)(Oughtred et al., 2021), Bioplex (Interactions with probability over 0.95)(Huttlin et al., 2021), human cellmap (Go et al., 2021), IntAct (only experimentally validated physical interactions)(Orchard et al., 2014), PINA2 (Cowley et al., 2012), and STRING (only with a STRING score > 0.9) databases (Szklarczyk et al., 2019). Number of citations per RTK were taken from gene2pubmed.gz file provided by NCBI at ftp://ftp.ncbi.nlm.nih.gov/gene/DATA/gene2pubmed.gz (May 2020). Domain annotations were mapped from PFam (El-Gebali et al., 2019), Reactome annotations from Uniprot to lowest pathway level mapping file available at Reactome (Fabregat et al., 2018). Gene ontology and CORUM (Giurgiu et al., 2019) annotations were taken from UniProt. GOCC annotations for CORUM complexes were taken from the CORUM database (Giurgiu et al., 2019). Known phosphosites, and kinases if available, were mapped from human protein reference database (Peri et al., 2003), PhosphoSitePlus (Hornbeck et al., 2015), phospho.ELM. (Dinkel et al., 2011), and a dataset from Sugiyama et al. (2019).

We checked expression status of our high-confidence interactors and our bait proteins against the Human Protein Atlas database version 20.1. (Uhlen et al. 2017) using RNA HPA cell line gene data that details the expression levels per gene in 69 different cell lines and against the RNA consensus tissue gene data that summarizes expression per gene in 62 tissues (downloaded 8.7.2021). In both cases ‘not expressed’ was judged to be a missing value or < 1 normalized expression (NX) value.

### Bioinformatic analyses

Enrichment values were calculated with an in-house python script using all identified proteins before any filtering steps were applied as the background set. Prey-Prey cross-correlation was calculated with in-house python script using scipy (Virtanen et al., 2020), and prey-prey associations from the correlation matrix were filtered based on q-value (< 0.01) calculated with scipy using FDR correction (Benjamini and Hochberg, 1995), and correlation value (> 0.7). Kinase domain sequence based clustering was done with clustal Omega (Madeira et al., 2019) using default settings and kinase domain sequences extracted from UniProt. Clustering of phosphotyrosine sites, and AP-MS and BioID data was performed in R with the seqinr and dendextend libraries. Clustering for heatmaps was performed in python using seaborn. Network figures were drawn with cytoscape 3.7 (Kohl et al., 2011). Fold change values for KD RTKs vs WT were calculated with an in-house python script using the WT kinase interactome as the background set. Random networks were generated by replacing HCIs in the RTK interactome with random proteins drawn from the background set of all identified proteins before any filtering steps were applied.

### Immunofluorescence confocal microscopy

The specific RTK expressing HEK293 cells were grown on glass coverslips. After 24 hours, cells were washed with PBS prior to fixation in 4% (wt/vol) paraformaldehyde (PFA) in PBS for 15 min at room temperature. Cells were then washed with PBS and permeabilized by 4 min of incubation in 0.1% (wt/vol) Triton X-100 in PBS. Bait proteins were detected with the anti-HA antibody (Thermo Fisher Scientific, Cat. No. 26183, dilution 1:1000 dilution), followed by Alexa Fluor488-conjugated secondary antibody (Thermo Fisher Scientific, A-11001, 1:1000 dilution). The nucleus was stained with DAPI (Sigma, Cat. No. D9542). Finally, coverslips were dried before mounting in Mowiol 4-88(Sigma, Cat. No. 81381). Prepared slides were analyzed using a confocal microscope (Leica TCS SP8 STED, Leica) with HC PL APO 93×/1.30 motCORR glycerol objective. Images were processed using ImageJ software (MacBiophotonics).

### Signal pathway analysis and luciferase assay

Cignal 45-Pathway Reporter Array (Qiagen, Cat. No. 336841) was used to monitor the corresponding signaling pathway activity. Briefly, 30 μl Opti-MEM containing dilute Attractene Transfection Reagent (Qiagen, Cat. No. 301005) was added to each well of the Cignal Finder Array plate coated with pre-formulated, transfection-ready reporter construct and test gene of interest construct, incubating at room temperature for 20 min. Subsequently, was added to DNA construct mixtures, 100 μl of HEK293 cell suspension containing 4 × 10^4^ cells in DMEM medium with 10% of fetal bovine serum was added to each well. After 24 h of transfection, the medium was changed to complete growth medium and further incubated for 24 h, followed by Dual-Luciferase Reporter Assay System that was performed according to the manufacturer’s protocol (Promega, Cat. No. E1960).

### Co-Immunoprecipitation

To validate the protein-protein interactions, HEK293 cells were co-transfected using Fugene 6 transfection reagent (Promega) with MAC-tag (600ng) and V5-tag (600ng) bait and prey constructs on 6-well cell culture plates with 5,00,000 cells per well. 24 hours post-transfection cells were rinsed with ice-cold 1X PBS and lysed with 1ml HENN lysis buffer per well (50mM HEPES pH8.0+ 5mM EDTA+ 150mM NaCl+ 50mM NaF+ 0.5% IGEPAL+ 1mM DTT+ 1mM PMSF+ 1.5mM Na3VO4 + 1X Protease inhibitor cocktail (Sigma-Aldrich)). Cell lysates were vortexed briefly and centrifuged (16,000 × g, 20min, 4°C) to remove cellular debris. 20 µl of Strep-Tactin® Sepharose® resin (IBA Lifesciences GmbH) was washed in a microcentrifuge tube twice with 200 µl HENN lysis buffer (4,000 × g, 1 min, 4°C). The clear lysate was collected and added to the washed Strep-Tactin® Sepharose® resin and incubated on a rotating wheel (60min, 4°C). After incubation, the samples were centrifuged (4,000 × g, 1 min, 4°C), and the supernatant was discarded. The pellet was washed three times with 1ml HENN lysis buffer (4,000 × g, 30 sec, 4°C). After the last wash, 60 µl of 2X Laemmli sample buffer was added directly to the beads and boiled at 95°C for 5 minutes. Samples were later used for immunodetection using western blot. For western blotting, immunoprecipitated proteins were detected with monoclonal mouse anti-V5 (Invitrogen) or mouse anti-HA.11 (BioLegend) primary antibodies and polyclonal goat anti-mouse HRP conjugated (GE Healthcare) secondary antibody. Signals were visualized by chemiluminescence using Amersham™ ECL™ Prime (Cytiva) for 5 min prior to imaging using iBright Imaging Systems (Thermo Fisher Scientific)

## Declaration

### Ethics approval and consent to participate

No patients or study participants were used in the study. Only commercially available existing cultured cell lines were used in this study.

### Consent for publication

No patient or study participants were used in the study.

### Conflict of Interests

The authors declare that they have no conflict of interest.

### Funding

This work was supported by the Academy of Finland and we thank Biocenter Finland/DDCB for financial support.

### Authors’ contributions

KS and MV designed the study. KS generated cell lines and performed AP-MS and BioID analyses. KS analyzed the AP-MS and BioID data. TÖ performed the IVK experiments. KS and TÖ analyzed the IVK data. XL performed the luciferase analysis and IF imaging. IC performed co-IP experiments. LG analyzed expression data from protein atlas. SK performed blotting experiments. KS and MV prepared the figures. KS, TÖ, and MV wrote the manuscript.

## Acknowledgements

We thank S. Miettinen for technical assistance and Professors Matthias Gstaiger, Aki Manninen and Kaisa Lehti for critical reading and comments on the manuscript. This study was supported by grants from the Academy of Finland (nos. 288475 and 294173), the Sigrid Jusélius Foundation, the Finnish Cancer Foundation, the University of Helsinki Three-year Research Grant, Biocentrum Helsinki, Biocentrum Finland, HiLIFE, Magnus Ehrnrooth Foundation and the Instrumentarium Research Foundation.

## Availability of data and material

Collected mass-spectrometry data is available at MassIVE with dataset ID MSV000087816, accessible with username “MSV000087816_reviewer” and password “rtk_varjosalo”. Plasmids are available from the corresponding author upon reasonable request.

